# Class I histone deacetylase HDA-3 is required for full maintenance of locomotor ability in *Caenorhabditis elegans*

**DOI:** 10.1101/794974

**Authors:** Kazuto Kawamura, Ichiro N. Maruyama

## Abstract

Locomotor ability declines with old age. A person’s capacity to maintain locomotor ability depends on genetic and environmental factors. Currently, the specific genetic factors that work to maintain locomotor ability are not well understood. Here we report the involvement of *hda-3*, encoding a class I histone deacetylase, as a specific genetic factor that contributes to the maintenance of locomotor ability in *C. elegans*. From a forward genetic approach, we identified a missense mutation in HDA-3 as the causative mutation for progressive decline in locomotor ability in one of the isolated strains. From transcriptome analysis, we found downregulated expression of two clusters of genes on Chromosome II and IV in this strain. Genes carrying CUB-like domains and genes carrying BATH domains were found on Chromosome II and IV, respectively. Knockdown of CUB-like genes, *K08D8.5* and *dod-17*, and BATH genes, *bath-1*, *bath-21* and *bath-24* led to a progressive decline in locomotor ability. Our study identifies specific genetic factors that work to maintain locomotor ability and reveals potential targets for delaying age-related locomotor decline.

## INTRODUCTION

Locomotor ability is a key determinant of quality of life in the elderly (Groessl *et al*. 2007). Age-related declines in locomotor ability are predictors of loss of independence, depressive symptoms, morbidity, and mortality (Trombetti *et al*. 2016). A combination of genetic and environmental factors contribute to how well a person can maintain locomotor ability during adulthood. Currently, the specific genetic factors that contribute to the maintenance of locomotor ability are largely unknown. A better understanding of the genetic factors that work to maintain locomotor ability may enable novel approaches to prevent or delay age-related declines in locomotor ability.

In order to identify genetic factors that regulate adult locomotor ability, we previously carried out a forward genetic screen for *C. elegans* mutants that show progressive declines in adult locomotor ability (Kawamura and Maruyama 2019). Characterization of one of the isolated strains led to the identification of a nonsense mutation in *elpc-2* and implicated the Elongator complex and tRNA modifications as factors that regulate locomotor healthspan in *C. elegans* (Kawamura and Maruyama 2019). Mutation in human ELP3, the catalytic subunit of the Elongator complex, has been linked to amyotrophic lateral sclerosis which suggests the evolutionarily conserved nature of the genetic factors that regulate locomotor healthspan (Simpson *et al*. 2009; Bento-Abreu *et al*. 2018).

In the present study, we analyzed another strain, *ix241*, to identify other genes that contribute to progressive decline in locomotor ability. In this strain, two notable mutations remained after four backcrosses: a splice site mutation in *dys-1* and a missense mutation in *hda-3* that leads to a glycine to glutamic acid substitution at the 271^st^ amino acid (G271E) in HDA-3. DYS-1 is the *C. elegans* ortholog of human Dystrophin, the causative gene that is mutated in Duchenne and Becker muscular dystrophies (Hoffman *et al*. 1987; Bessou *et al*. 1998). HDA-3 is a *C. elegans* ortholog of human class I histone deacetylases HDAC1–3 (Shi and Mello 1998). Surprisingly, mutation in *hda-3*, but not *dys-1*, contributed to progressive decline in locomotor ability during adulthood. Downstream of the G271E mutation in HDA-3, specific genes carrying CUB-like domains and genes carrying BATH domains are transcriptionally repressed. Proper induction of CUB-like and BATH genes are required for full maintenance of locomotor ability during adulthood.

## MATERIALS AND METHODS

### Strains

*C. elegans* Bristol N2 strain was used as the wild type strain. Worms were cultivated at 20°C on Nematode Growth Media (NGM) agar plates with *Escherichia coli* strain OP50 as a food source (Brenner 1974). All strains used in this study are listed in Table S1.

### Sanger sequencing

The target genomic region was amplified using PCR and purified using Wizard SV Gel and PCR Clean-Up System (Promega, Madison, WI). The DNA sequence of the PCR fragment was determined using cycle sequencing with BigDye v3.1 reagents (Applied Biosystems, Foster City, CA). Sequencing products were purified by EtOH/EDTA precipitation. Sequencing was performed by capillary sequencing using ABI3100 (Applied Biosystems). Primers used for Sanger sequencing are listed in Table S2.

### Whole-genome DNA sequencing

*C. elegans* DNA was sequenced using the MiSeq next-generation sequencing system (Illumina, San Diego, CA) as previously described (Kawamura and Maruyama 2019). Libraries for sequencing were prepared with Illumina TruSeq Library Prep Kit. Sequenced reads were mapped using BWA software (Li and Durbin 2009). Mapped read files were converted to bam format, then to pileup format with Samtools (Li *et al*. 2009). Variant detection was carried out using VarScan and SnpEff (Blankenberg *et al*. 2010; Cingolani *et al*. 2012; Giardine *et al*. 2005; Goecks *et al*. 2010; Koboldt *et al*. 2009). Mutation frequencies were calculated and visualized using CloudMap (Minevich *et al*. 2012).

### Measurements of maximum speed and travel distance

“Synchronized egg-laying” was used to raise a batch of worms of similar age. Five adult day 1 worms were placed onto an NGM plate with food, and allowed to lay eggs for 3 h. When the offspring reached adult day 1, 15 worms were randomly picked onto a 6 cm NGM plate without bacteria. After the worms moved away from the initial location with residual food, worms were again moved onto a different NGM plate without bacteria. The maximum speed and travel distance of worms were measured on the first, third, and fifth days of adulthood as previously described (Kawamura and Maruyama 2019). R was used to make plots (Team 2015).

### CRISPR-Cas9 genome editing

Targeted mutagenesis was carried out using CRISPR-Cas9 genome editing with single-stranded oligodeoxynucleotide (ssODN) donors as previously described (Dokshin *et al*. 2018). First, a ribonucleoprotein complex was created by mixing together 0.5 μL of 10 μg/μL Cas9 protein, 5.0 μL of 0.4 μg/μL of tracrRNA, and 2.8 μL of 0.4 μg/μL of crRNA (Target-specific sequence: 5’-CCGAUUCACUGGCAGGAGAU-3’) and incubating at 37°C for 10 min. Following incubation, 2.2 μL of 1 μg/μL ssODN, 2.0 μL of 400 ng/μL pRF4::*rol-6(su1006)* co-injection marker, and 7.5 μL of nuclease free water was added to the mixture. This mixture was then injected into the gonad of worms subject to genomic editing. F1 offspring that showed the roller phenotype were singled onto individual plates, and allowed to lay eggs. Editing of the target sequence was checked by single worm PCR of the F1 worm, followed by Sanger sequencing. ssODN sequences are listed in Table S3.

### RNA sequencing

Worms were synchronized by placing ten adult day 1 worms onto an NGM plate with food, and allowed to lay eggs for 3 h. At the L4 stage, worms were collected and washed with M9 buffer and placed on 9 cm NGM plates with 25 μM floxuridine (FUDR). On the third day of adulthood, worms were collected with M9 buffer and RNA was extracted. Worms were homogenized using Micro Smash MS-100R (Tomy Seiko, Tokyo, Japan). RNA was extracted by the phenol-choloroform method using Trizol reagent (Thermo Fisher Scientific). Sequencing was performed on the HiSeq platform (Illumina). For bioinformatics analysis, reads were aligned using STAR (Dobin *et al*. 2013), sorting and marking duplicates were done by Picard, and read counting was done by Featurecounts (Liao *et al*. 2014). EdgeR (Robinson *et al*. 2009) and R (Team 2015) were used to create figures to visualize differential gene expression.

### Quantitative PCR (qPCR)

RNA was extracted by the phenol-choloroform method using Trizol reagent (Thermo Fisher Scientific). cDNA was synthesized using SuperScript III with oligo-dT primers (Thermo Fisher Scientific). qPCR was carried out with Luna Universal qPCR Master Mix (New England Biolabs, Ipswich, MA) using StepOnePlus (Thermo Fisher Scientific).

### RNA interference

The Ahringer RNAi library was used to reduce the expression of target genes (*F55G11.8*, *K08D8.5*, *K10D11.1*, *F59H6.8*, *F59H6.9*, *B0047.3*). Frozen stocks of the RNAi bacteria were streaked onto agar plates containing 100 μg/mL ampicillin. Single colonies of the RNAi bacteria were cultured for 8 h at 37°C with vigorous shaking in lysogeny broth (LB) containing 100 μg/mL ampicillin. 100 μL of bacterial culture was spread on NGM plates with 50 μg/mL ampicillin and 1.0 mM isopropyl β-D-1-thiogalactopyranoside (IPTG). Plates with RNAi bacteria were dried overnight with the lid on at room temperature (25°C). Adult wild-type worms were placed on RNAi plates and allowed to lay eggs for 3 h. The locomotor ability of the offspring was tested from the first day of adulthood. For mock control, an empty vector L4440 was used.

### Statistics

All results are expressed as means with error bars representing a 95% confidence interval. For pairwise comparisons, Student’s *t* test was used with Excel 2010 (Microsoft). For multiple comparisons to a control, one-way ANOVA was followed with Dunnett’s post hoc test using R (Team 2015). For multiple comparisons, one-way ANOVA was followed with Tukey’s Honest Significant Difference test using R (Team 2015). Statistical significance was set at \**P* < 0.05; \*\**P* < 0.01; \*\*\**P* < 0.001.

### Data availability

All isolated strains are available upon request. DNA and RNA sequencing results are available on NCBI sequence read archive PRJNA530333.

## RESULTS

### Identification of novel *dys-1(ix259)* mutant allele in *ix241* strain

Previously, we carried out a forward genetic screen for *C. elegans* mutants with a shortened adult locomotor healthspan (Kawamura and Maruyama 2019). One of the isolated strains was the *ix241* strain which shows a slight developmental deficit in locomotor ability and a progressive decline in locomotor ability during adulthood (Kawamura and Maruyama 2019). The *ix241* strain retained an exaggerated head bending phenotype after four backcrosses, suggesting that the phenotype may be linked with progressive decline in locomotor ability (Figure 1A). Exaggerated head bending has previously been observed by numerous independent research groups in mutants with loss-of-function mutations to *dys-1*, the *C. elegans* ortholog of human Dystrophin, and to components of the Dystrophin associated protein complex (DAPC) (Fig. S1A) (Oh *et al*. 2012; Kim *et al*. 2004, 2009; Grisoni *et al*. 2003; Zhou and Chen 2011; Bessou *et al*. 1998).

**Figure 1.**
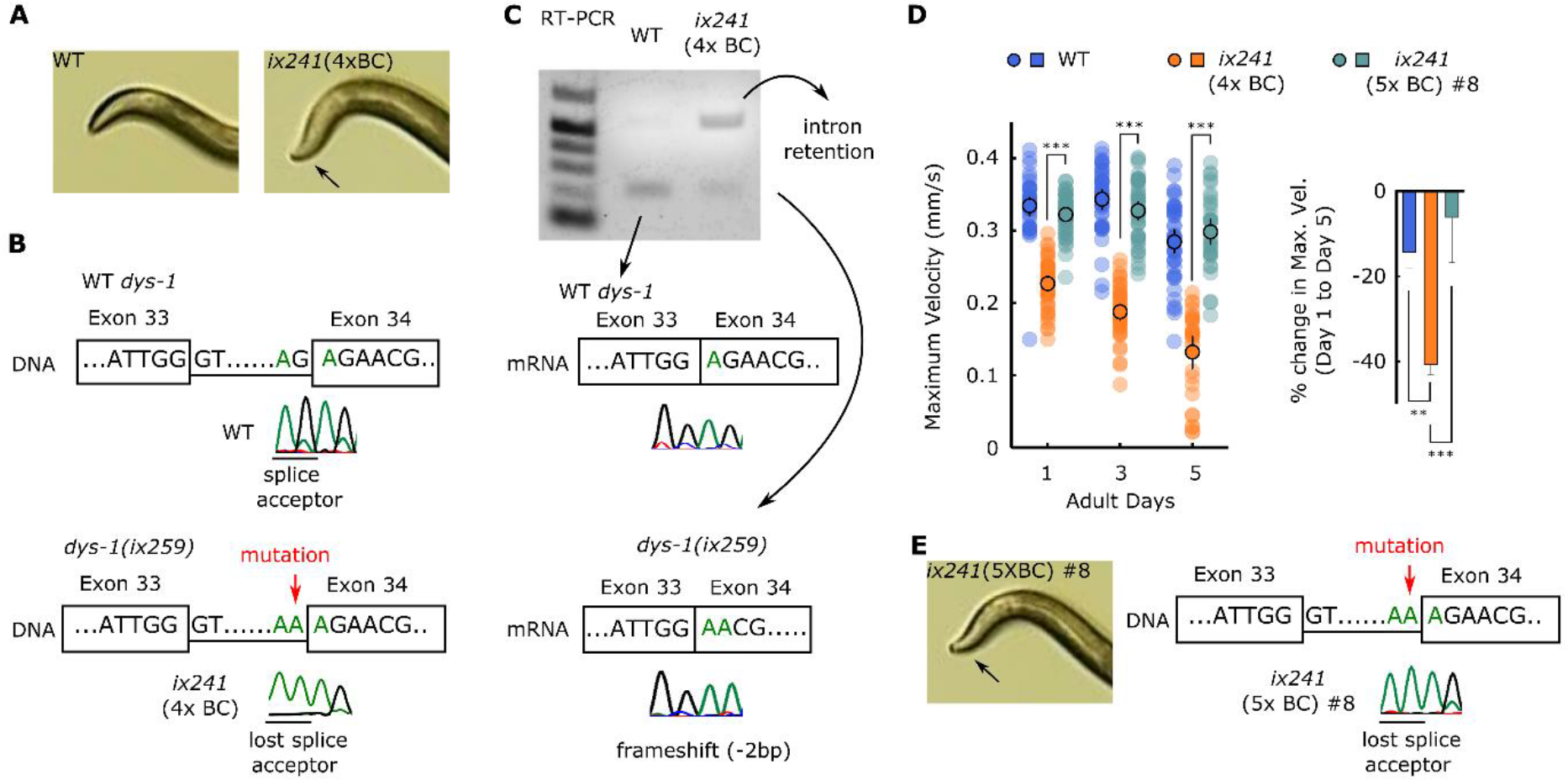
Novel *dys-1(ix259)* loss-of-function mutation allele is present in *ix241* strain but does not cause progressive decline in locomotor ability.

(A) Photos of head curvature during forward crawling in WT and *ix241*(4x BC) worms. *ix241*(4x BC) worms show exaggerated head bending (indicated by arrow). (B) (*Top*) DNA sequence of *dys-1*(*ix259*) mutation site from WT worms and (*Bottom*) *ix241*(4x BC) worms. (C) (Top) cDNA sequence of *dys-1*(*ix259*) mutation site from WT and (Bottom) *ix241*(4x BC) worms. (D) (*Left*) Maximum velocities of WT, *ix241*(4x BC), and *ix241*(5x BC) #8 worms. n=30–45 worms per strain for each day (10–15 worms from 3 biological replicate plates). (*Right*) Percent change in maximum velocity of WT, *ix241*(4x BC), and *ix241*(5x BC) #8 worms on adult day 5 compared to adult day 1. n=3 biological replicate plates. \*\*\**P* < 0.001; \**P* < 0.05. (E) (*Left*) Photo of head curvature during forward crawling in *ix241*(5x BC) #8 worms. (*Right*) DNA sequence of *dys-1*(*ix259*) mutation site in *ix241*(5x BC) #8 worms.

Whole genome sequencing of backcrossed *ix241* strains that show progressive declines in locomotor ability revealed a splice site mutation in *dys-1* prior to the 34^th^ exon, which was confirmed by Sanger sequencing (Figure 1B, Table S4). We refer to this mutation site as *dys-1(ix259)*, since later we found that this mutation site is not involved in the progressive decline in locomotor ability. Reverse-transcriptase PCR using primers that flank the *dys-1(ix259)* splice site mutation indicated that intron retention occurs in the majority of *dys-1* mRNA in the *ix241* strain (Figure 1C). A small proportion of *dys-1* transcripts are spliced using an adjacent splice site, but results in a 2 bp frameshift (Figure 1C). Intron retention or the 2 bp frameshift would likely lead to nonsense-mediated mRNA decay of *dys-1* transcripts.

### *dys-1* mutations do not cause progressive decline in locomotor ability in *C. elegans*

In humans, Dystrophin mutations cause progressive weakness of muscles (Hoffman *et al*. 1987). Therefore, we hypothesized that the *dys-1(ix259)* splice site mutation may be the causative mutation site for the progressive decline in locomotor ability in the *ix241* strain. However, after the fifth backcross we isolated *ix241*(5x BC) #8, a strain that carries the *dys-1(ix259)* mutation but does not show a progressive decline in locomotor ability as measured by maximum velocity and travel distance (Figure 1D, 1E, Fig. S1B, S1C). The *ix241*(5x BC) #8 strain shows the exaggerated head bending phenotype observed in *dys-1* mutants (Figure 1E). The phenotype of the *ix241*(5x BC) #8 worms raised the possibility that *dys-1(ix259)* does not lead to progressive decline in locomotor ability in the *ix241* strain.

We wondered whether other *dys-1* mutants show a progressive decline in locomotor ability. There are two available *dys-1* mutants from the *Caenorhabditis* Genetics Center: BZ33 strain carrying *dys-1(eg33)* and LS292 strain carrying *dys-1(cx18)*. Both *dys-1(eg33)* and *dys-1(cx18)* mutant alleles are nonsense mutations. Interestingly, *dys-1(eg33)* mutant worms show a progressive decline in locomotor ability while *dys-1(cx18)* mutant worms do not show a progressive decline in locomotor ability from the first to fifth days of adulthood (Figure 2A, Fig. S2A). Similar to our findings, *dys-1(eg33)* worms, but not *dys-1(cx18)* worms were found to have significantly weaker adult muscle strength compared to wild-type worms (Hewitt *et al*. 2018). The discrepancy was attributed to a difference in the *dys-1* mutation allele. However, in light of the newly isolated *dys-1(ix259)* mutant worms which do not show a progressive decline in locomotor ability, we hypothesized that the progressive decline in adult locomotor ability in the BZ33 strain may not be caused by the *dys-1(eg33)* mutation.

**Figure 2.**
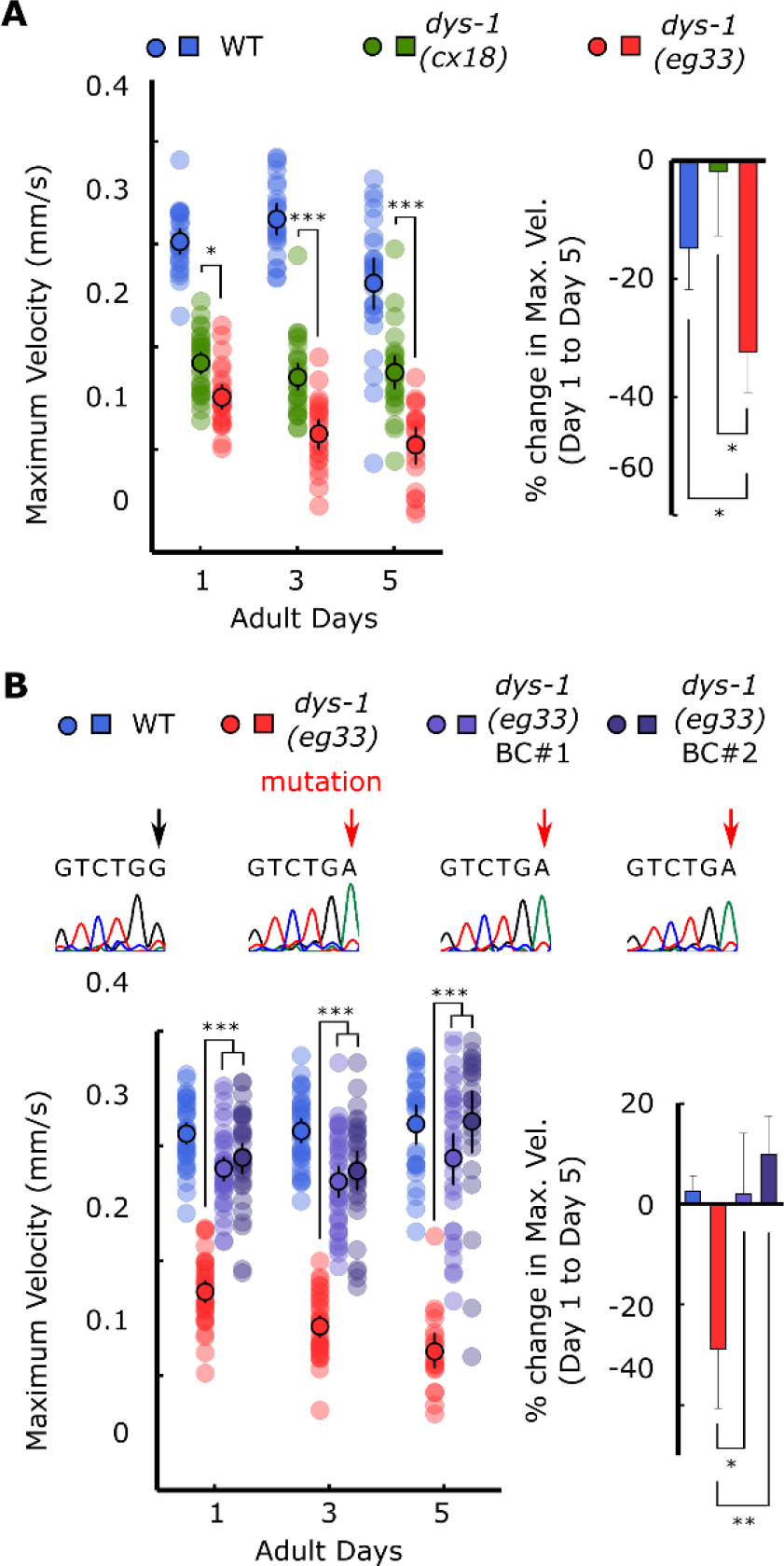
*dys-1(eg33)* mutation does not cause progressive decline in locomotor ability from first to fifth day of adulthood.

(A) (*Left*) Maximum velocities of WT, *dys-1(cx18)*, and *dys-1(eg33)* worms. (*Right*) Percent change in maximum velocity of WT, *dys-1(cx18)*, and *dys-1(eg33)* worms on adult day 5 compared to adult day 1. (B) (*Top*) DNA sequences of *dys-1(eg33)* mutation site in WT, *dys-1(eg33)*, and two independent backcrossed lines, *dys-1(eg33)* BC #1, and *dys-1(eg33)* BC #2 worms. (*Left*) Maximum velocities of WT, *dys-1(eg33)*, *dys-1(eg33)* BC #1 and *dys-1(eg33)* BC#2 worms. (*Right*) Percent change in maximum velocity of WT, *dys-1(eg33)*, *dys-1(eg33)* BC #1, and *dys-1(eg33)* BC #2 worms on adult day 5 compared to adult day 1. For maximum velocity measurements, n=30–45 worms per strain for each day (10–15 worms from 3 biological replicate plates). For percent change in maximum velocity graphs, n=3 biological replicate plates. \*\*\**P* < 0.001; \*\**P* < 0.01; \**P* < 0.05.

In order to test whether the progressive decline in locomotor ability in the BZ33 strain is caused by a mutation aside from *dys-1(eg33)*, we backcrossed the BZ33 strain based on the exaggerated head bending phenotype (Fig. S2B). After one backcross, we were able to isolate two strains that carry the *dys-1(eg33)* mutation and show the exaggerated head bending, but do not show progressive decline in locomotor ability from the first to fifth days of adulthood (Figure 2B, Fig. S2C). This suggests that loss-of-function mutations in *dys-1* does not cause progressive decline in locomotor ability from the first to fifth days of adulthood in *C. elegans*. A mutation site aside from *dys-1(eg33)* likely causes progressive decline in locomotor ability in the BZ33 strain.

### *hda-3* mutation causes progressive decline in locomotor ability in *ix241* strain

In order to identify the causative mutation site that leads to progressive decline in locomotor ability in the *ix241* strain, we carried out whole genome sequencing in strains that showed and did not show the progressive decline in locomotor ability after backcrossing. We identified the mutations that were shared among the genomes of *ix241* backcrossed strains that showed the progressive decline in adult locomotor function, and subtracted the shared mutations among *ix241* backcrossed strains that did not show the progressive decline in adult locomotor function. A peak of mutations remained on Chromosome I (Figure 3A, 3B, Table S5).

**Figure 3.**
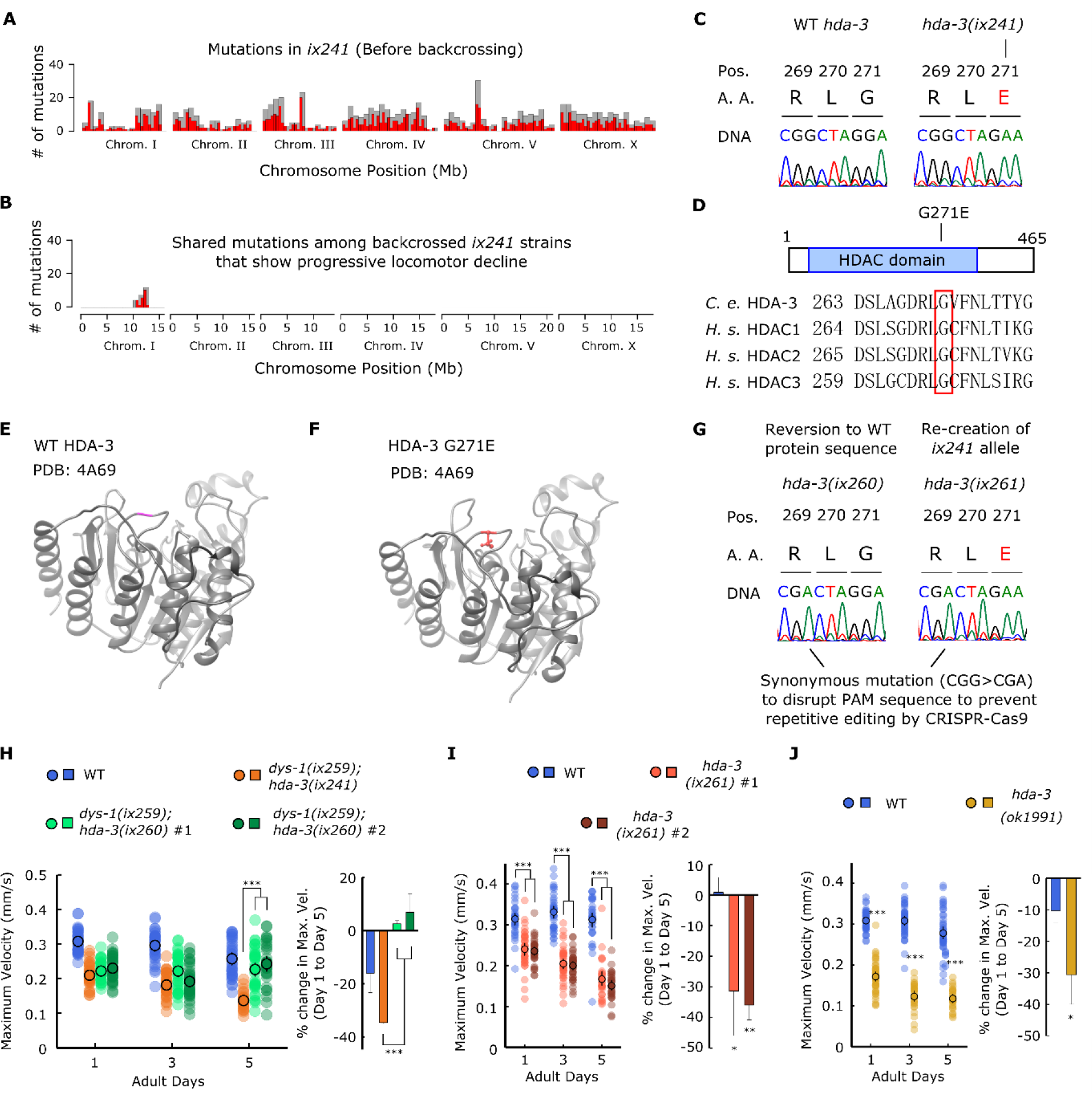
HDA-3 G271E missense mutation leads to progressive decline in locomotor ability.

(A) Mutation frequency along each chromosome for the *ix241* strain before backcrossing. Red bars indicate 0.5-Mb bins and grey bars indicate 1.0-Mb bins. (B) Mutation frequency along each chromosome for remaining mutations after subtracting mutations found in *ix241* backcrossed strains that did not show a progressive decline in locomotor ability from mutations found in *ix241* backcrossed strains that did show a progressive decline in locomotor ability. (C) Effect of *ix241* mutation on DNA sequence and amino acid sequence. (D) (*Top*) Depiction of *hda-3(ix241)* mutation site in HDA-3 protein. (*Bottom*) Alignment of amino acid sequences centered around G271E mutation site in *C. elegans* HDA-3, *H. sapiens* HDAC1, HDAC2 and HDAC3. (E) Structural modeling of *C. elegans* WT HDA-3 based on PDB: 4A69 from *H. Sapiens* HDAC3. (F) Structural modeling of *C. elegans* HDA-3 G271E based on PDB: 4A69 from *H. Sapiens* HDAC3. Mutated glutamic acid residue is shown in red. (G) (Left) Sequence of *hda-3(ix260)* allele which is the same amino acid sequence as wild type. (Right) Sequence of *hda-3(ix261)* allele which is the same amino acid sequence as the *hda-3(ix241)* allele. Both sequences carry a synonymous mutation site to disrupt the PAM sequence to prevent repetitive editing by CRISPR-Cas9. (H) (*Left*) Maximum velocities of WT, *dys-1(ix259);hda-3(ix241)* worms, *dys-1(ix259);hda-3(ix260)* #1 and *dys-1(ix259);hda-3(ix260)* #2 worms. (*Right*) Percent change in maximum velocity of WT, *dys-1(ix259);hda-3(ix241)* worms, *dys-1(ix259);hda-3(ix260)* #1 and *dys-1(ix259);hda-3(ix260)* #2 worms on adult day 5 compared to adult day 1. (I) (*Left*) Maximum velocities of WT, *hda-3(ix261)* #1, *hda-3(ix261)* #2 worms. (*Right*) Percent change in maximum velocity of WT, *hda-3(ix261)* #1, *hda-3(ix261)* #2 worms on adult day 5 compared to adult day 1. (J) (*Left*) Maximum velocities of WT and *hda-3*(*ok1991)* worms. (*Right*) Percent change in maximum velocity of WT and *hda-3(ok1991)* worms on adult day 5 compared to adult day 1. For maximum velocity measurements, n=30–45 worms per strain for each day (10–15 worms from 3 biological replicate plates). For percent change in maximum velocity graphs, n=3 biological replicate plates. \*\*\**P* < 0.001; \*\**P* < 0.01; \**P* < 0.05.

We identified a list of candidate mutation sites, which included *hda-3(ix241)* that would cause a G271E missense mutation in HDA-3 (Figure 3C, 3D). HDA-3 is an ortholog of human class I histone deacetylases, HDAC1–3 (Shi and Mello 1998). The G271 residue is evolutionarily conserved from *C. elegans* to humans and is located in the variable loop region (Figure 3D, 3E, 3F). The variable loop region is suggested to play a role in substrate recognition and binding to the HDAC cofactors zinc and inositol phosphate (Watson *et al*. 2012; Schuetz *et al*. 2008) (Figure 3E, 3F).

The effect of the *hda-3(ix241)* mutation was tested by two strategies. In the first strategy, CRISPR-Cas9 genome editing was used to revert the *hda-3(ix241)* mutation in the *ix241*(4x BC) strain back to the WT sequence (Fig. 3G). In order to prevent repetitive editing, a synonymous mutation was introduced that would disrupt the protospacer adjacent motif (PAM) sequence, 5 bp upstream of the editing site (Fig. 3G). We refer to this reverted allele as *hda-3(ix260)*, which has the same HDA-3 amino acid sequence as WT HDA-3 (Fig. 3G). In the second strategy, the HDA-3 G271E mutation was introduced into the WT N2 background using CRISPR-Cas9 genome editing. Again, a synonymous mutation was introduced that would disrupt the PAM sequence, 5 bp upstream of the editing site (Fig. 3G). We refer to this mutation allele as *hda-3(ix261)*, which causes the same HDA-3 G271E mutation as the *hda-3(ix241)* mutation (Fig. 3G). Strains carrying *hda-3(ix261)* were backcrossed twice.

The reversion of the *hda-3(ix241)* mutation in the *ix241*(4x BC) strain to *hda-3(ix260)* rescued the progressive decline in locomotor function (Figure 3H, Fig. S3A). This result indicated that the *hda-3(ix241)* mutation is necessary for the progressive decline in locomotor function in the *ix241* strain. Introduction of the G271E mutation in the N2 WT strain in *hda-3(ix261)* strains led to progressive declines in locomotor ability (Figure 3I). This result indicated that the HDA-3 G271E mutation alone is sufficient to cause progressive decline in locomotor ability. In addition, an independently isolated *hda-3(ok1991)* deletion strain showed progressive decline in locomotor ability (Figure 3J, Fig. S3C). Proper functioning of HDA-3 is likely to be required for full maintenance of locomotor ability during adulthood.

### Expression of specific CUB-like and BATH genes are dysregulated in *hda-3* mutant strains

In order to identify gene expression changes that occur in the *ix241*(4x BC) strain, transcriptome analysis was carried out. In comparison to wild-type worms, *ix241*(4x BC) worms had 64 transcripts that were significantly upregulated and 47 transcripts that were significantly downregulated (Figure 4A). In comparison to *ix241*(5x BC) #8 worms, *ix241*(4x BC) worms had 27 transcripts that were significantly upregulated and 25 transcripts that were significantly downregulated (Figure 4A). Twenty-two transcripts were commonly upregulated in the *ix241*(4x BC) strain compared to wild type and the *ix241*(5x BC) #8.Thirteen transcripts were commonly downregulated in the *ix241*(4x BC) strain compared to wild type and the *ix241*(5x BC) #8 (Figure 4B). Gene ontology enrichment analysis indicated that transcripts involved in the immune response were significantly enriched in both upregulated and downregulated transcripts (Fig. S4A, B).

**Figure 4.**
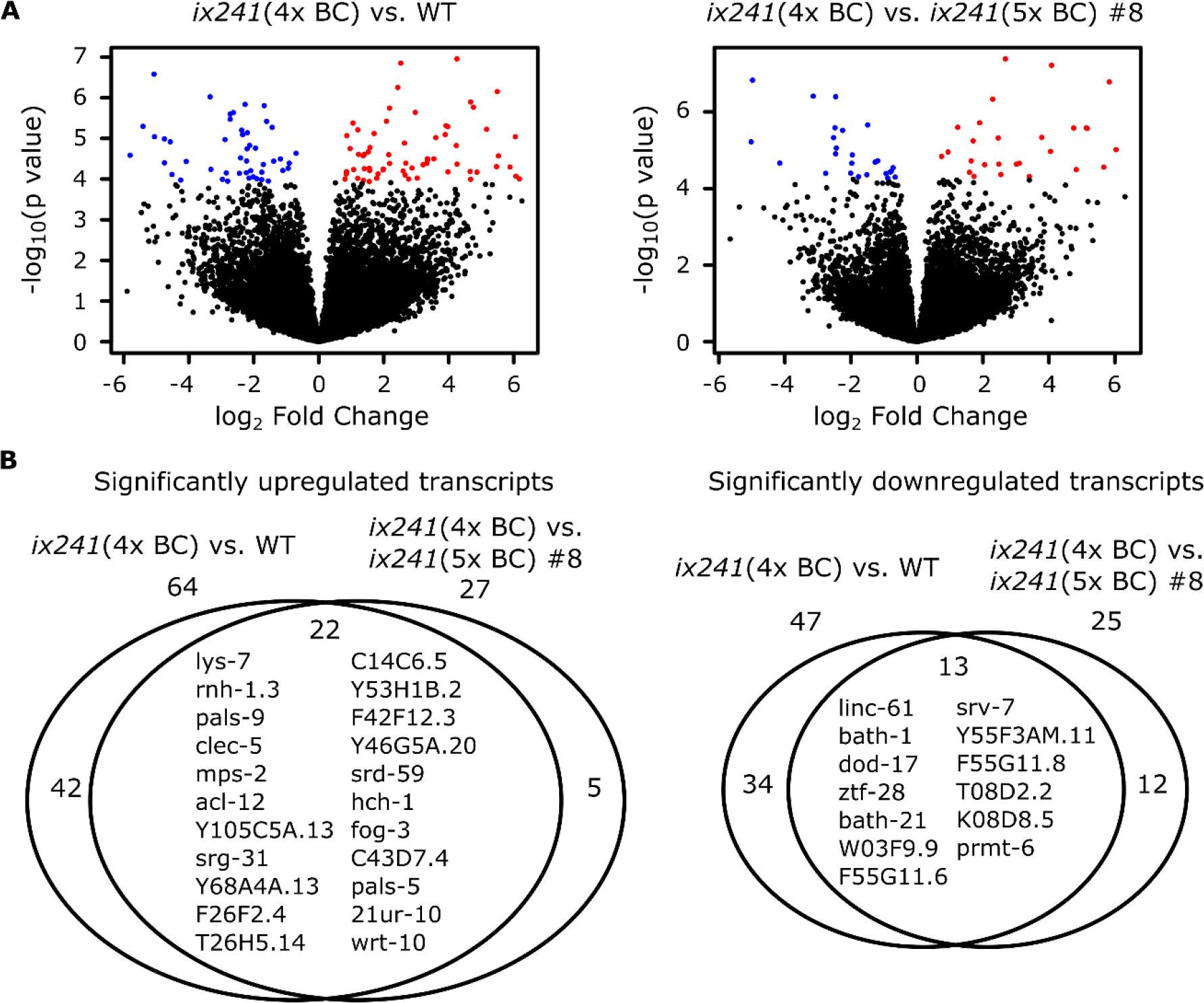
Gene expression is dysregulated in *hda-3* mutant.

(A) (*Left*) Volcano plot of differential expression of transcripts from *ix241*(4x BC) vs. WT worms. (*Right*) Volcano plot of differential expression of transcripts from *ix241*(4x BC) vs. *ix241*(5x BC) #8 worms. Blue points indicate downregulated genes and red points indicate upregulated genes with *p* value < 0.0001. (B) (*Left*) Venn diagram of number of significantly upregulated transcripts with *p* <0.0001 in *ix241*(4x BC) vs. WT worms and *ix241*(4x BC) vs. *ix241*(5x BC) #8 worms. (*Right*) Venn diagram of number of significantly upregulated and downregulated transcripts with *p* <0.0001 in *ix241*(4x BC) vs. WT and *ix241*(4x BC) vs. *ix241*(5x BC) #8 worms. Names of commonly upregulated or downregulated genes are indicated within the Venn diagram.

Among the downregulated transcripts, we noticed that multiple gene transcripts were downregulated within two narrow regions of the genome. One of the downregulated regions is on Chromosome II, where BATH domain carrying proteasome-related genes *bath-1*, *bath-21*, and *bath-24* are located (Figure 5A). The other downregulated region was on Chromosome IV where CUB-like domain carrying innate immune response genes *dod-17*, *F55G11.6*, *F55G11.8*, *K08D8.5* are located (Figure 5A). All genes showed high levels of expression except for *F55G11.6*, which showed very low expression levels in WT and *ix241*(5xBC) #8. Downregulation of CUB-like and BATH genes were also seen in *hda-3(ix261)* mutant worms (Figure 5B). In the *hda-3(ok1991)* deletion mutant, *bath-1*, *bath-21*, *bath-24*, and *F55G11.8* were downregulated while *dod-17* and *K08D8.5* remained unchanged (Figure 5C).

**Figure 5.**
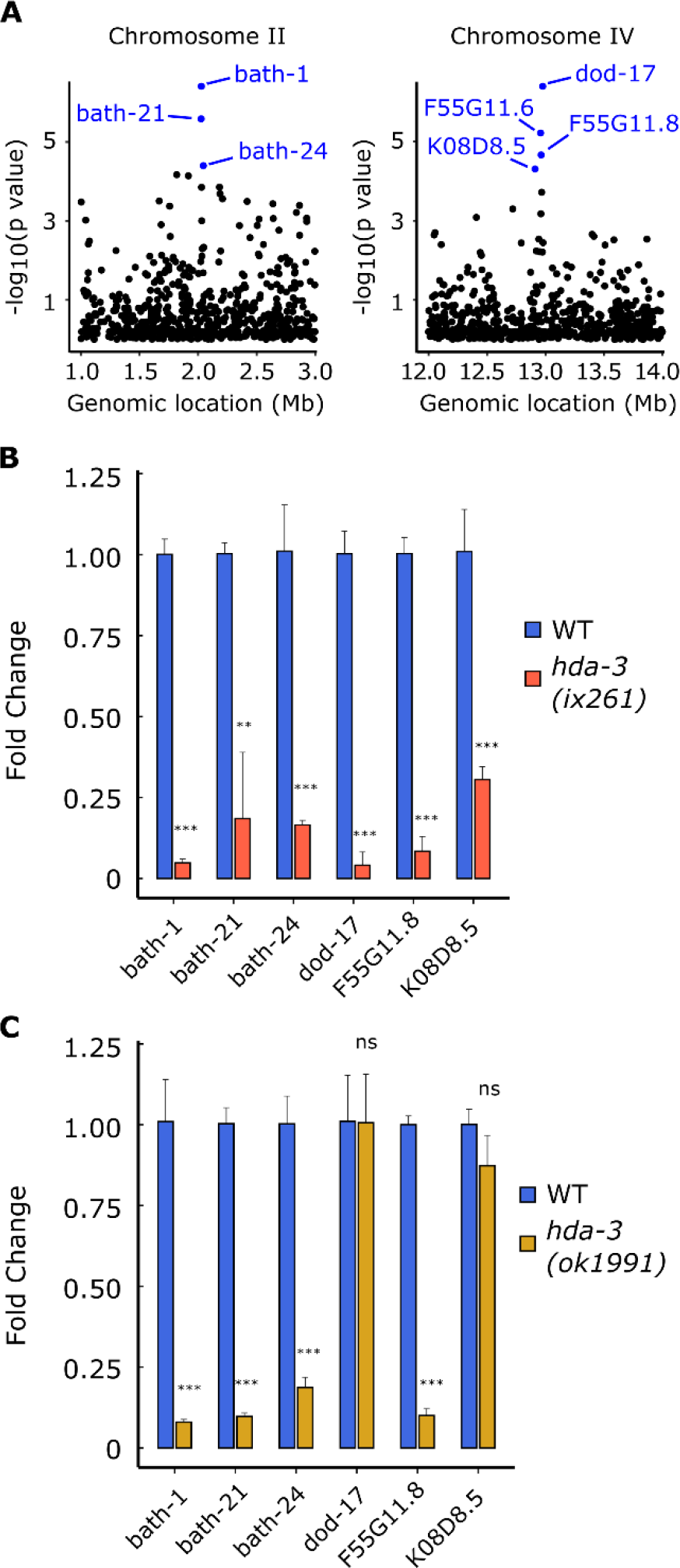
Specific BATH and CUB-like genes on chromosome II and IV are downregulated from HDA-3 G271E mutation.

(A)(*Left*) Genomic location of strongly downregulated gene transcripts on Chromosome II. (*Right*) Genomic location of strongly downregulated gene transcripts on Chromosome IV. (B) Fold change in CUB-like and BATH genes in *hda-3(ix261)* worms compared to WT, as measured by qPCR. (C) Fold change in CUB-like and BATH genes in *hda-3(ok1991)* worms compared to WT, as measured by qPCR. \*\*\**P* < 0.001; \*\**P* < 0.01.

### Induction of CUB-like and BATH genes are required for full maintenance of locomotor ability

We tested whether the downregulation of the CUB-like and BATH genes contribute to the progressive decline in locomotor function. We knocked down the CUB-like and BATH genes in wild-type worms and measured their locomotor ability for seven days. Knockdown of five out of the six genes led to a significant decline in locomotor ability on the fifth day of adulthood (Figure 6A, 6B, Fig. S5A, S5B). We observed significant declines in locomotor ability as compared from the first to fifth day of adulthood in five out of the six tested genes (Figure 6A, 6B, Fig. S5A, S5B).

**Figure 6.**
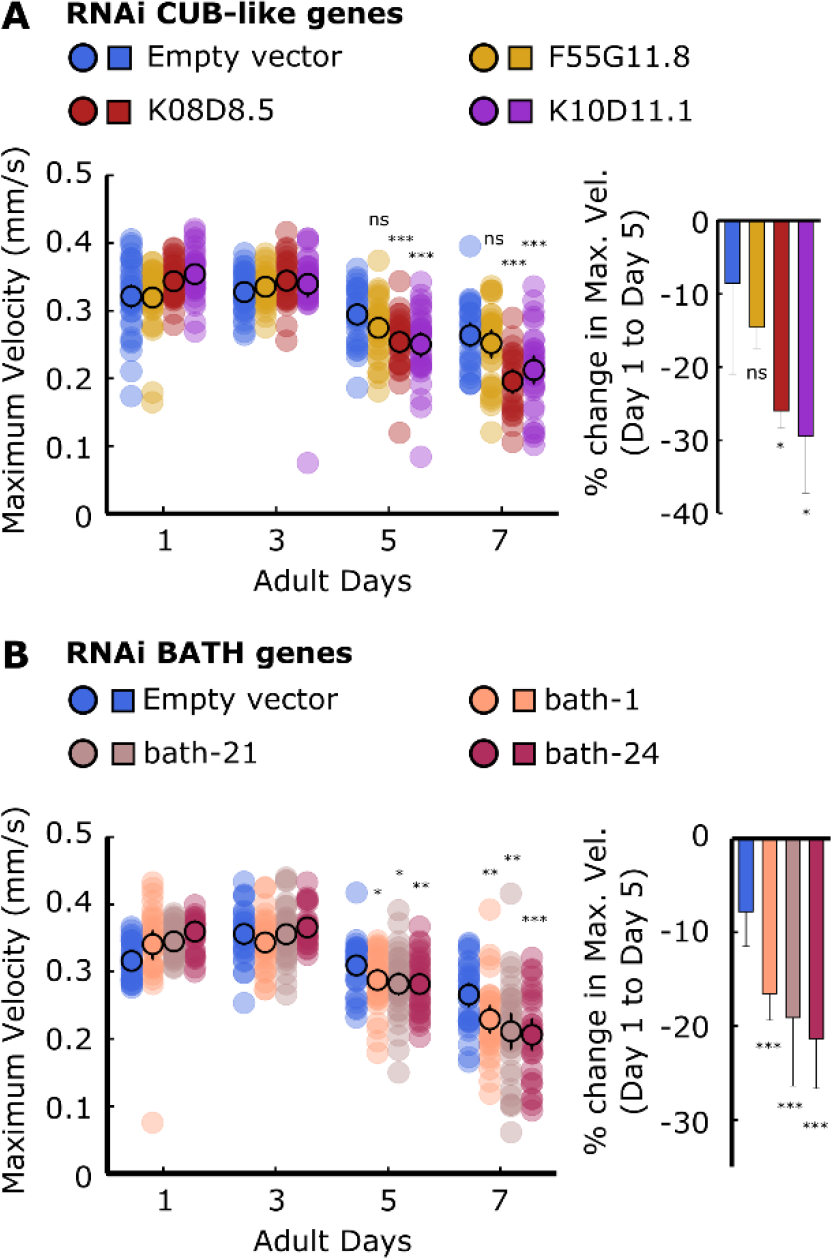
RNAi of certain CUB-like and BATH genes cause progressive decline in locomotor ability.

(A) (*Left*) Maximum velocities of worms raised on RNAi plates containing empty vector, *F55G11.8, K08D8.5, K10D11.1*. (*Right*) Percent change in maximum velocity of worms raised on RNAi plates containing empty vector, *F55G11.8, K08D8.5, K10D11.1* on adult day 5 compared to adult day 1. (B) (*Left*) Maximum velocities of worms raised on RNAi plates containing empty vector, *bath-1, bath-21, bath-24*. (*Right*) Percent change in maximum velocity of worms raised on RNAi plates containing empty vector, *bath-1, bath-21, bath-24* on adult day 5 compared to adult day 1. For maximum velocity measurements, n=30–45 worms per strain for each day (10–15 worms from 3 biological replicate plates). For percent change in maximum velocity graphs, n=3 biological replicate plates. \*\*\**P* < 0.001; \*\**P* < 0.01; \**P* < 0.05.

## DISCUSSION

In this study, we found that proper HDA-3 function is required for the full maintenance of locomotor ability in *C. elegans*. In the *ix241* strain, progressive decline in locomotor ability is caused by the *hda-3(ix241)* mutant allele which leads to a G271E substitution in HDA-3. In *hda-3* mutants carrying the G271E mutation, we observed specific downregulation of gene clusters located on chromosome II and IV. The downregulated cluster of genes on chromosome II carry a CUB-like domain, and downregulated genes on chromosome IV carry a BATH domain. Knockdown of several of the most significantly downregulated CUB-like and BATH genes leads to a progressive decline in locomotor ability. This study indicates the importance of proper HDA-3 functioning and induction of genes carrying CUB-like or BATH domains for the maintenance of adult locomotor ability.

HDA-3 is a histone deacetylase that can affect the transcriptional expression of many downstream genes (Struhl 1998). Generally, histone acetylation is positively associated with transcriptional activation and deacetylation is associated with transcriptional repression (Eberharter and Becker 2002). However, some studies have found that HDACs are involved in both repression and activation (Wang *et al*. 2002; Nusinzon and Horvath 2005). Our transcriptomics results show a similar number of upregulated and downregulated genes in the *hda-3(ix241)* mutant strain, and support the notion that HDACs have dual roles in transcriptional repression and activation.

Transcriptome analysis and quantitative PCR of mutant strains carrying the HDA-3 G271E mutation indicated two regions in the genome that are transcriptionally repressed on Chromosome II and IV. The Chromosome II region carried multiple genes that contain BATH domains, which are suggested to work as part of the immunoproteasome to target foreign proteins for degradation (Thomas 2006). The Chromosome IV region carried multiple genes that contain CUB-like domains, which are implicated in innate immune function (Bork and Beckmann 1993). Knockdown of the most significantly repressed CUB-like and BATH genes caused progressive decline in locomotor function in *C. elegans*, indicating that these genes may be functional targets of HDA-3 to maintain locomotor ability in *C. elegans*. Expression of genes carrying CUB-like domains and genes carrying BATH domains may be important for the maintenance of locomotor ability. Interestingly, the structural properties of both the BATH domain and CUB-like domain are characterized by beta sandwiches which were first characterized in immunoglobulins. The *ix241* strain may enable further exploration of the link between the innate immune system and neuromuscular maintenance.

The G271E mutation occurs at an evolutionarily conserved residue. The same residue is present in human HDAC1–3. This residue is likely a critical residue for proper HDAC function and may enable novel approaches for the inhibition of HDACs. The G271 amino acid is located on one of the four variable loop regions which is implicated in substrate recognition (Schapira 2011). The G271 amino acid is in close proximity to R269, which is an evolutionarily conserved amino acid that mediates the interaction between human HDAC3 and its coactivator, inositol phosphate (Watson *et al*. 2012; Millard *et al*. 2013). D263 is also a nearby amino acid which is predicted to mediate the interaction between HDAC family proteins with the cofactor zinc (Schuetz *et al*. 2008). The G271 amino acid site may provide a novel location for drug targets to alter the activity of class I histone deacetylases.

The role of class I HDACs during the aging process has been difficult to study, as HDAC1, HDAC2, HDAC3, and HDAC8 have been found to play important roles during development in vertebrates (Haberland *et al*. 2009). For example, HDAC3 knockout mice die during embryonic development (Montgomery *et al*. 2008). Since *C. elegans* can tolerate the loss of *hda-3* during development, the HDA-3 G271E mutants and *hda-3(ok1991)* deletion mutant may be valuable tools to study the role of a specific class I histone deacetylase during aging.

During the process of identifying the causative mutation site of the *ix241* strain, we found that *dys-1* loss-of-function mutations do not cause a progressive decline in adult locomotor ability. Sufficient backcrossing showed that the progressive decline in locomotor ability does not segregate perfectly with the *dys-1* mutation site in the *ix241* strain and the BZ33 strain carrying the *dys-1(eg33)* mutant allele. This came as a surprise since DYS-1 is the ortholog of human Dystrophin, the causative mutation for Duchenne and Becker muscular dystrophies (Hoffman *et al*. 1987). Our findings suggest the use of caution when interpreting the role of *dys-1* in the maintenance of muscle strength or locomotor ability.

These findings should not preclude the use of *C. elegans dys-1* mutants to study potential mechanisms of Duchenne muscular dystrophy. Genetic and molecular interactions of Dystrophin are highly conserved in *C. elegans*. Dystrobrevin and syntrophin, which are components of the Dystrophin-associated protein complex, have *C. elegans* orthologs which interact with with *C. elegans* DYS-1 (Grisoni *et al*. 2003; Oh *et al*. 2012). One major advantage of *C. elegans* Dystrophin mutants is the presence of the exaggerated head bending phenotype, which can be readily observed under the microscope (Oh *et al*. 2012; Kim *et al*. 2004, 2009; Grisoni *et al*. 2003; Zhou and Chen 2011; Bessou *et al*. 1998). Future drug screenings and genetic manipulations that suppress the head bending phenotype in the *dys-1* mutants may be a promising avenue to identify modifiers of *dys-1* loss-of-function.

In a previous study, we identified a nonsense mutation in *elpc-2* that leads to progressive decline in locomotor ability (Kawamura and Maruyama 2019). The role of *elpc-2* as part of the Elongator complex implicates the role of tRNA modifications for the maintenance of proteostasis and adult locomotor ability. In this study, we identify the G271E mutation in HDA-3 and its role in transcriptional regulation of CUB-like and BATH genes for the maintenance of adult locomotor ability. Together, these mutants provide insights into the mechanisms that contribute to the maintenance of adult locomotor ability. Future studies of mutants that show progressive declines in locomotor ability may provide further insights into the genetic programs that work to maintain our locomotor healthspan.

## ACKNOWLEDGEMENTS

We thank H. Goto, M. Kanda, M. Kawamitsu, S. Yamasaki, N. Arakaki and other DNA sequencing section members for technical assistance with DNA and RNA sequencing. We are grateful to T. Murayama and E. Saita for technical support and advice. We thank D. Van Vactor, B. Kuhn, and members of the Maruyama unit for helpful discussions regarding this work. We are grateful to H. Ohtaki for administrative support. We thank the *Caenorhabditis* Genetics Center, which is funded by NIH Office of Research Infrastructure Programs (P40 OD010440), for providing worm strains. This work was partly supported by Okinawa Institute of Science and Technology Graduate University, and K. K. was supported by Japan Society for the Promotion of Science KAKENHI (Grant 16J06404).

---

**Figure S1.**
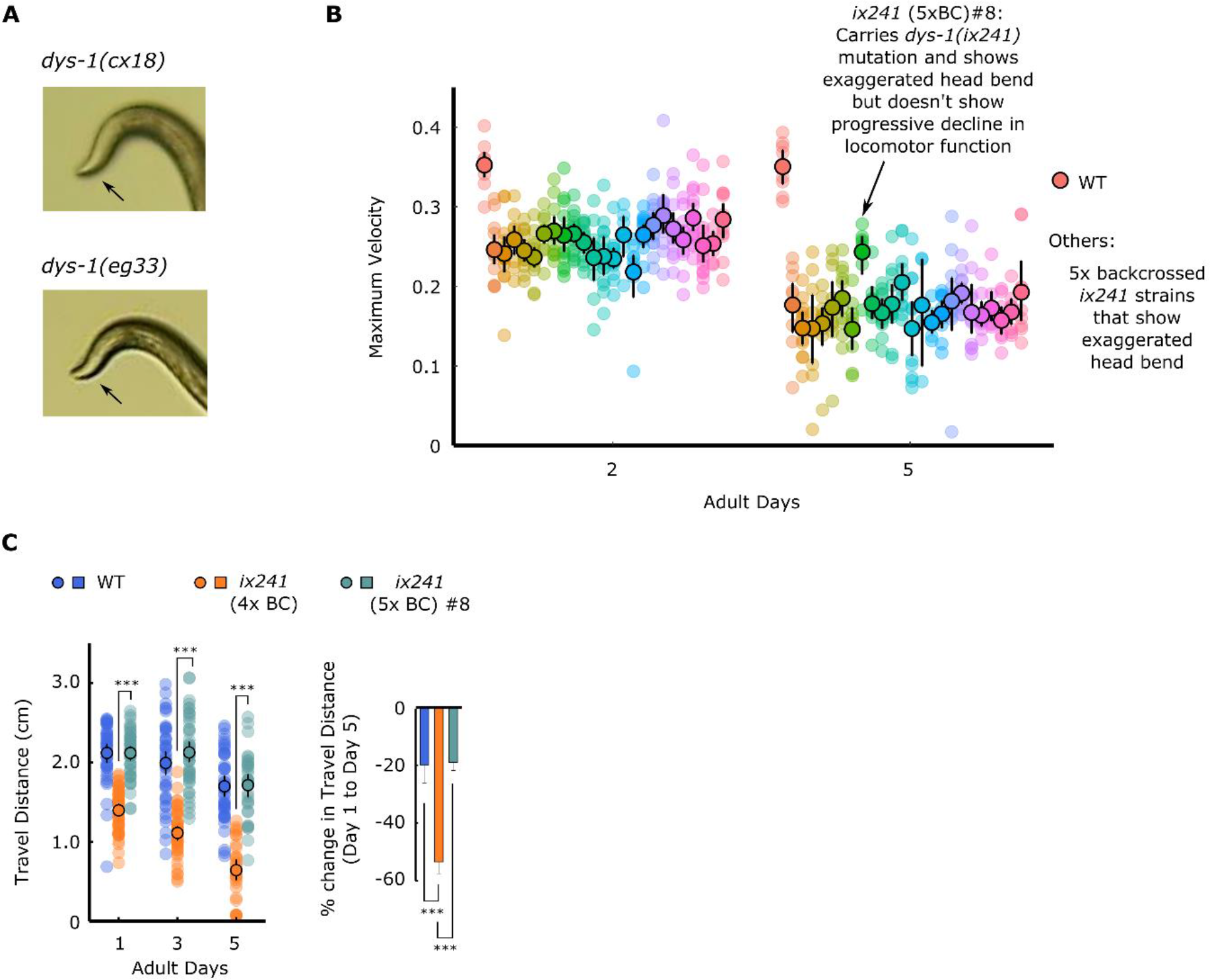
*ix241* (5xBC) #8 strain carries *dys-1(ix259)* mutation but does not show progressive decline in locomotor ability.

(A) Photos of head curvature during forward crawling in WT and *ix241*(4x BC) worms. Exaggerated head bending indicated by arrows. (B) Maximum velocities of 24 strains that show the exaggerated head bending phenotype after the fifth backcross on the second and fifth days of adulthood. n=10–15 worms per strain. (C) (*Left*) Travel distances of WT, *ix241*(4x BC), and *ix241*(5xBC) #8 worms. n=30–45 worms per strain for each day (10–15 worms from 3 biological replicate plates). (*Right*) Percent change in travel distance of WT, *ix241*(4x BC), and *ix241*(5xBC) #8 worms. n=3 biological replicate plates. \*\*\**P* < 0.001.

**Figure S2.**
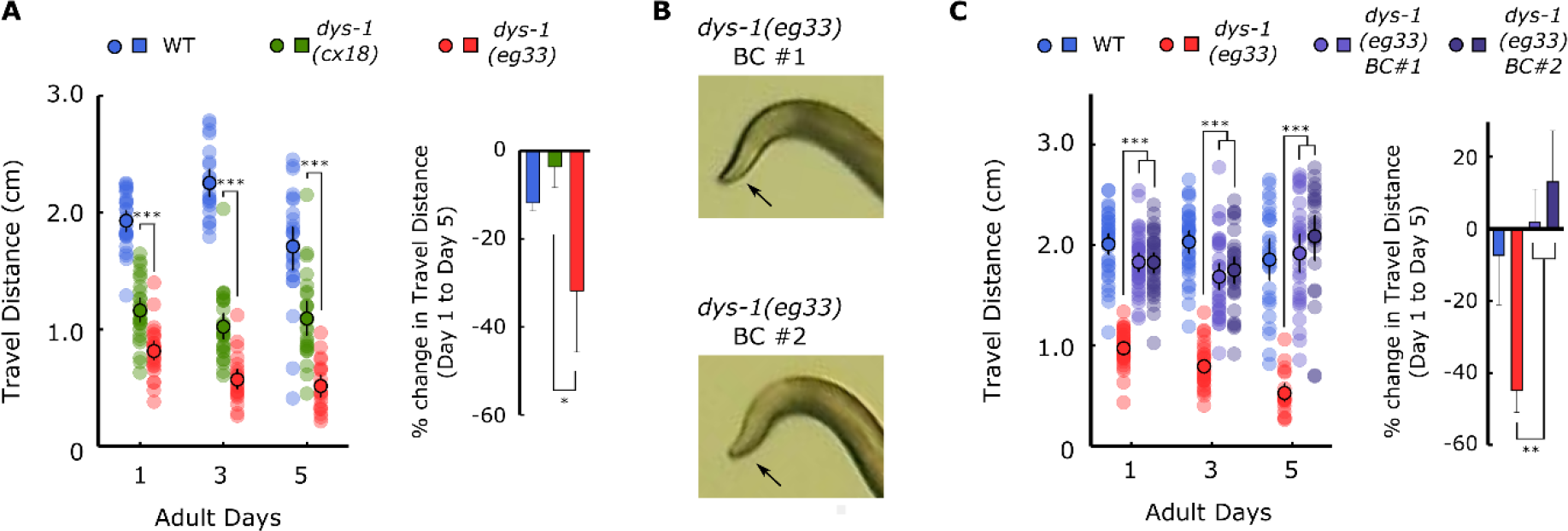
*dys-1(eg33)* mutation does not cause progressive decline in locomotor ability.

(A) (*Left*) Travel distances of WT, *dys-1(cx18)*, and *dys-1(eg33)* worms on adult days 1, 3, and 5. (*Right*) Percent change in travel distance of WT, *dys-1(cx18)*, and *dys-1(eg33)* worms on adult day 5 compared to adult day 1. (B) Photos of head curvature during forward crawling in *dys-1(eg33)* BC #1 and *dys-1(eg33)* BC #2 worms. Arrows indicate exaggerated head bending. (C) (*Left*) Travel distances of WT, *dys-1(eg33)*, *dys-1(eg33)* BC #1, and *dys-1(eg33)* BC #2 worms on adult days 1, 3, and 5. (*Right*) Percent change in travel distance of WT, *dys-1(eg33)*, *dys-1(eg33)* BC #1, and *dys-1(eg33)* BC #2 worms on adult day 5 compared to adult day 1. For travel distance measurements, n=30–45 worms per strain for each day (10–15 worms from 3 biological replicate plates). For percent change in travel distance graphs, n=3 biological replicate plates. \*\*\**P* < 0.001; \*\**P* < 0.01; \**P* < 0.05.

**Figure S3.**
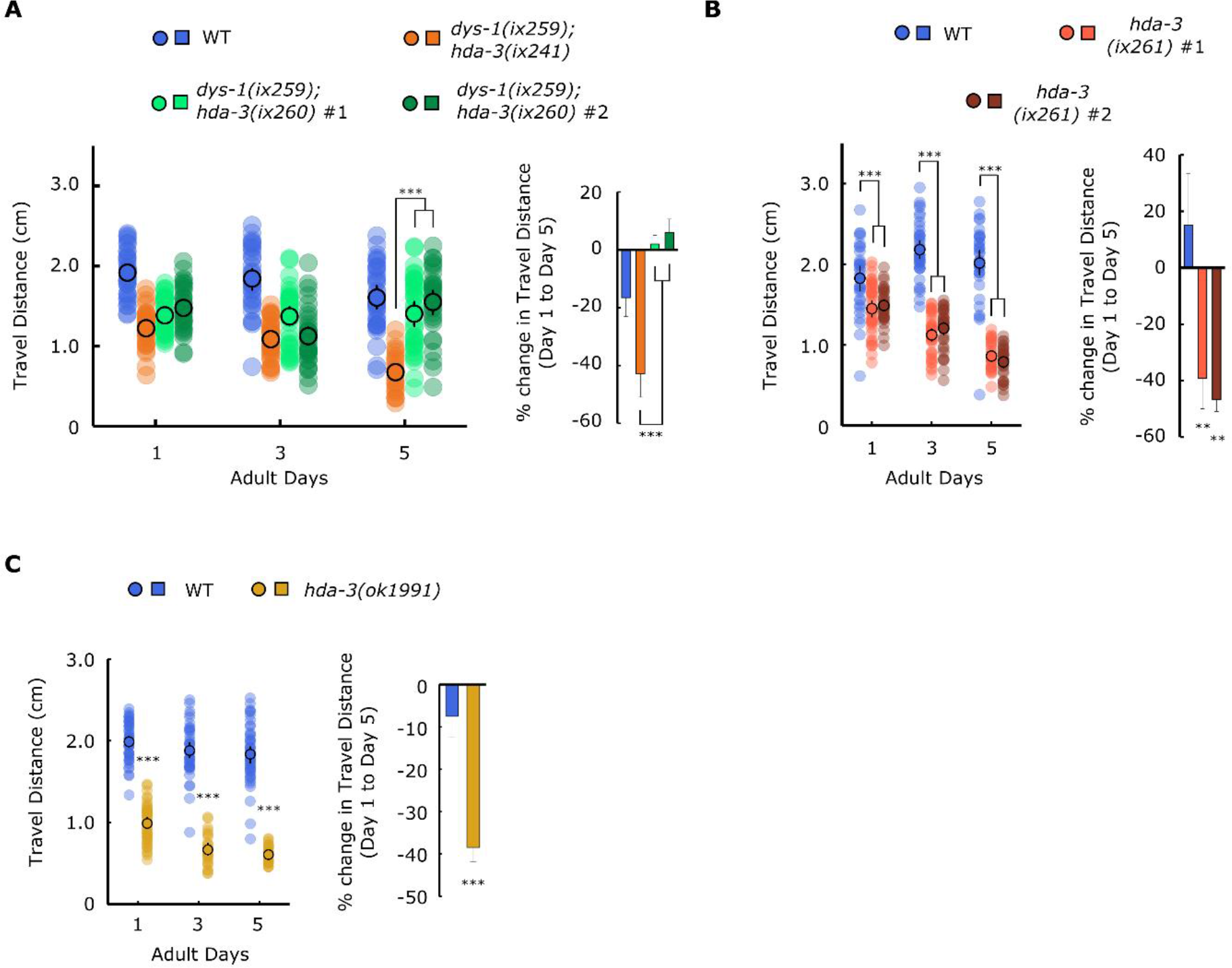
*hda-3* mutation causes progressive decline in locomotor ability.

(A) (*Left*) Travel distances of WT, *dys-1(ix259);hda-3(ix241)*, *dys-1(ix259);hda-3(ix260)* #1 and *dys-1(ix259);hda-3(ix260)* #2 worms on adult days 1, 3, and 5. (*Right*) Percent change in travel distance of WT, *dys-1(ix259);hda-3(ix241)*, *dys-1(ix259);hda-3(ix260)* #1 and *dys-1(ix259);hda-3(ix260)* #2 worms on adult day 5 compared to adult day 1. (B) (*Left*) Travel distances of WT, *hda-3(ix261)* #1, and *hda-3(ix261)* #2 worms on adult days 1, 3, and 5. (*Right*) Percent change in travel distance of WT, *hda-3(ix261)* #1, and *hda-3(ix261)* #2 worms on adult day 5 compared to adult day 1. (C) (*Left*) Travel distances of WT and *hda-3(ok1991)* worms on adult days 1, 3, and 5. (*Right*) Percent change in travel distance of WT and *hda-3(ok1991)* worms on adult day 5 compared to adult day 1. For travel distance measurements, n=30–45 worms per strain for each day (10–15 worms from 3 biological replicate plates). For percent change in travel distance graphs, n=3 biological replicate plates. \*\*\**P* < 0.001; \*\**P* < 0.01.

**Figure S4.**
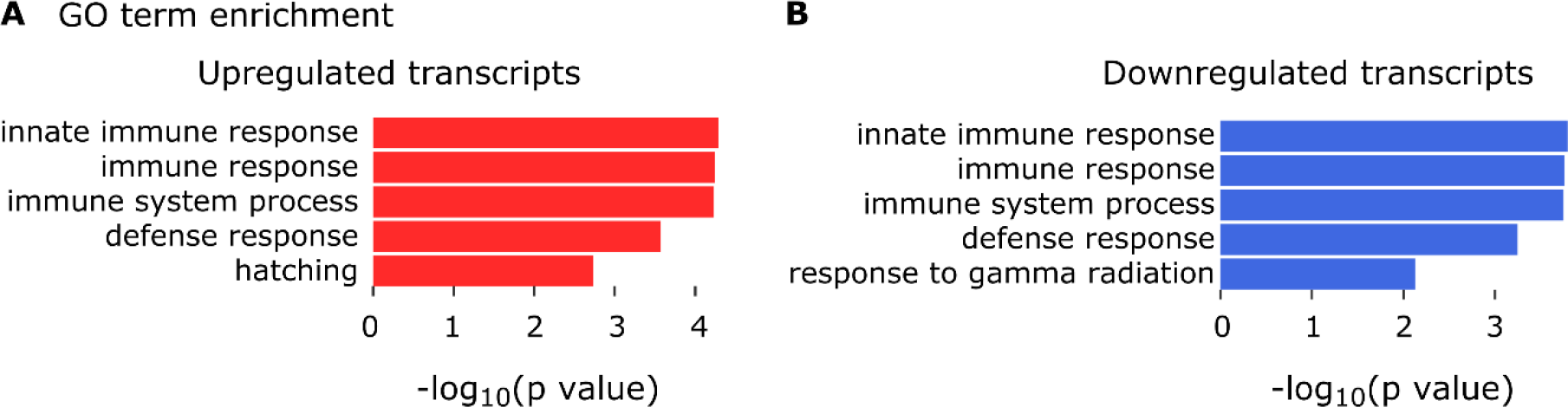
Dysregulated genes are enriched for those involved in immune function.

(A) Enriched GO terms among genes upregulated in *ix241*(4x BC) strain versus WT and *ix241* (5x BC) #8. (B) Enriched GO terms among genes downregulated in *ix241*(4x BC) strain versus WT and *ix241* (5x BC) #8.

**Figure S5.**
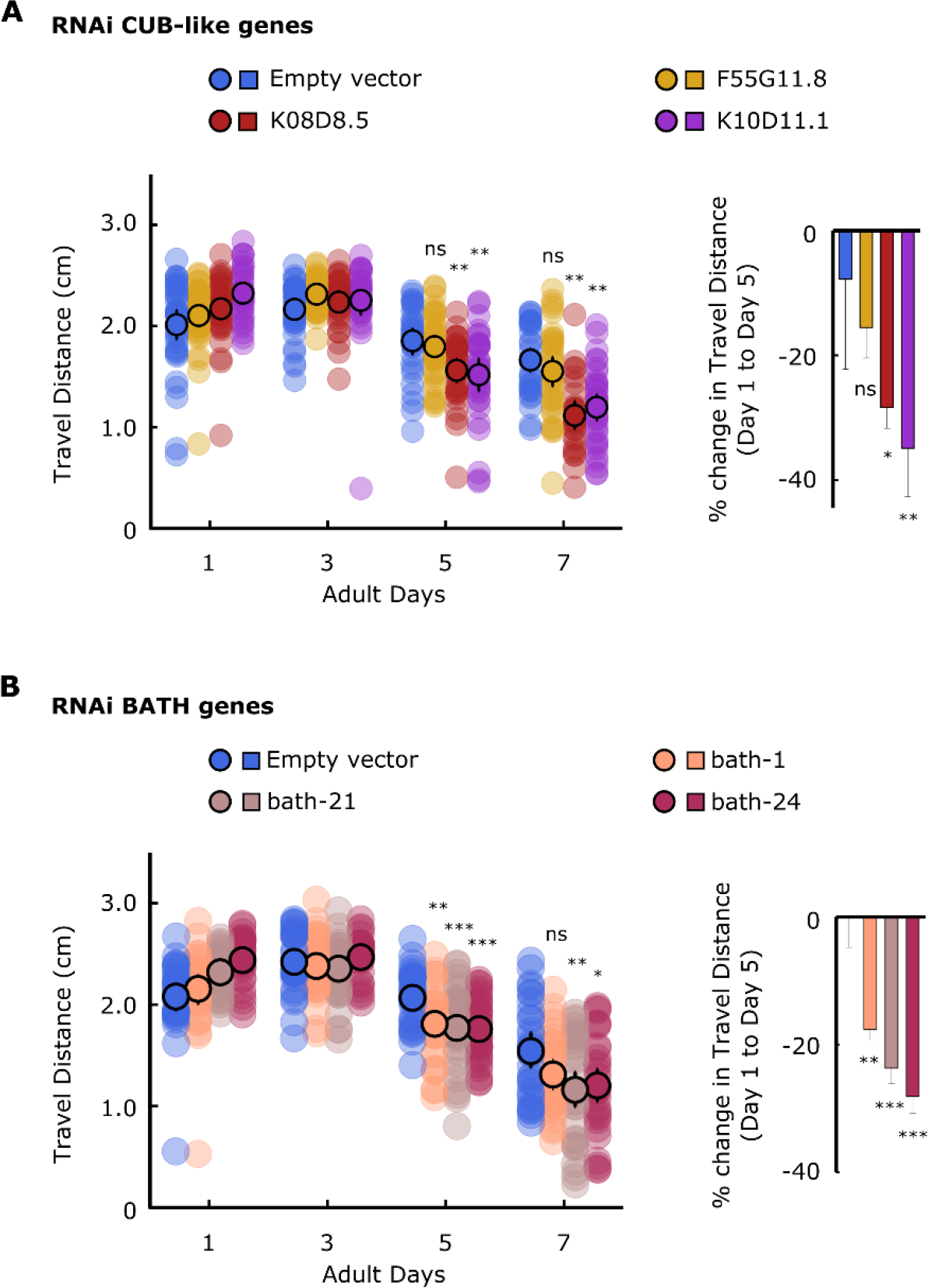
Knockdown of CUB-like and BATH genes lead to progressive decline in locomotor ability.

(A) (*Left*) Travel distances of worms raised on RNAi plates containing empty vector, *F55G11.8, K08D8.5, K10D11.1*. (*Right*) Percent change in travel distance of worms raised on RNAi plates containing empty vector, *F55G11.8, K08D8.5, K10D11.1* on adult day 5 compared to adult day 1. (B) (*Left*) Travel distances of worms raised on RNAi plates containing empty vector, *bath-1, bath-21, bath-24*. (*Right*) Percent change in travel distance of worms raised on RNAi plates containing empty vector, *bath-1, bath-21, bath-24* on adult day 5 compared to adult day 1. For travel distance measurements, n=30–45 worms per strain for each day (10–15 worms from 3 biological replicate plates). For percent change in travel distance graphs, n=3 biological replicate plates. \*\*\**P* < 0.001; \*\**P* < 0.01; \**P* < 0.05.

**Table S1.** List of strains used in this study.

| Strain | Genotype | Obtained from |
| --- | --- | --- |
| OF1262 | *dys-1(ix259) I; hda-3(ix241)* | Isolated in previous study (Kawamura and Maruyama, 2019);<br>Also referred to as *ix241* |
| OF1263 | *dys-1(ix259) I; hda-3(ix241) I* (4x backcrossed) | Isolated in previous study (Kawamura and Maruyama, 2019);<br>Also referred to as *ix241* (4x BC) |
| OF1350 | *dys-1(ix259) I* (5x backcrossed) | This study; Also referred to as *ix241* (5x BC) #8 |
| LS292 | *dys-1(cx18)* | CGC |
| BZ33 | *dys-1(eg33)* | CGC |
| OF1351 | *dys-1(eg33)* (1x backcrossed) #1 | This study |
| OF1352 | *dys-1(eg33)* (1x backcrossed) #2 | This study |
| OF1353 | *dys-1(ix259) I; hda-3(ix260)* #1 | This study |
| OF1354 | *dys-1(ix259) I; hda-3(ix260)* #2 | This study |
| OF1355 | *hda-3(ix261) I* (2x backcrossed) #1 | This study |
| OF1356 | *hda-3(ix261) I* (2x backcrossed) #2 | This study |
| RB1618 | *hda-3(ok1991) I* | CGC |

**Table S2.**
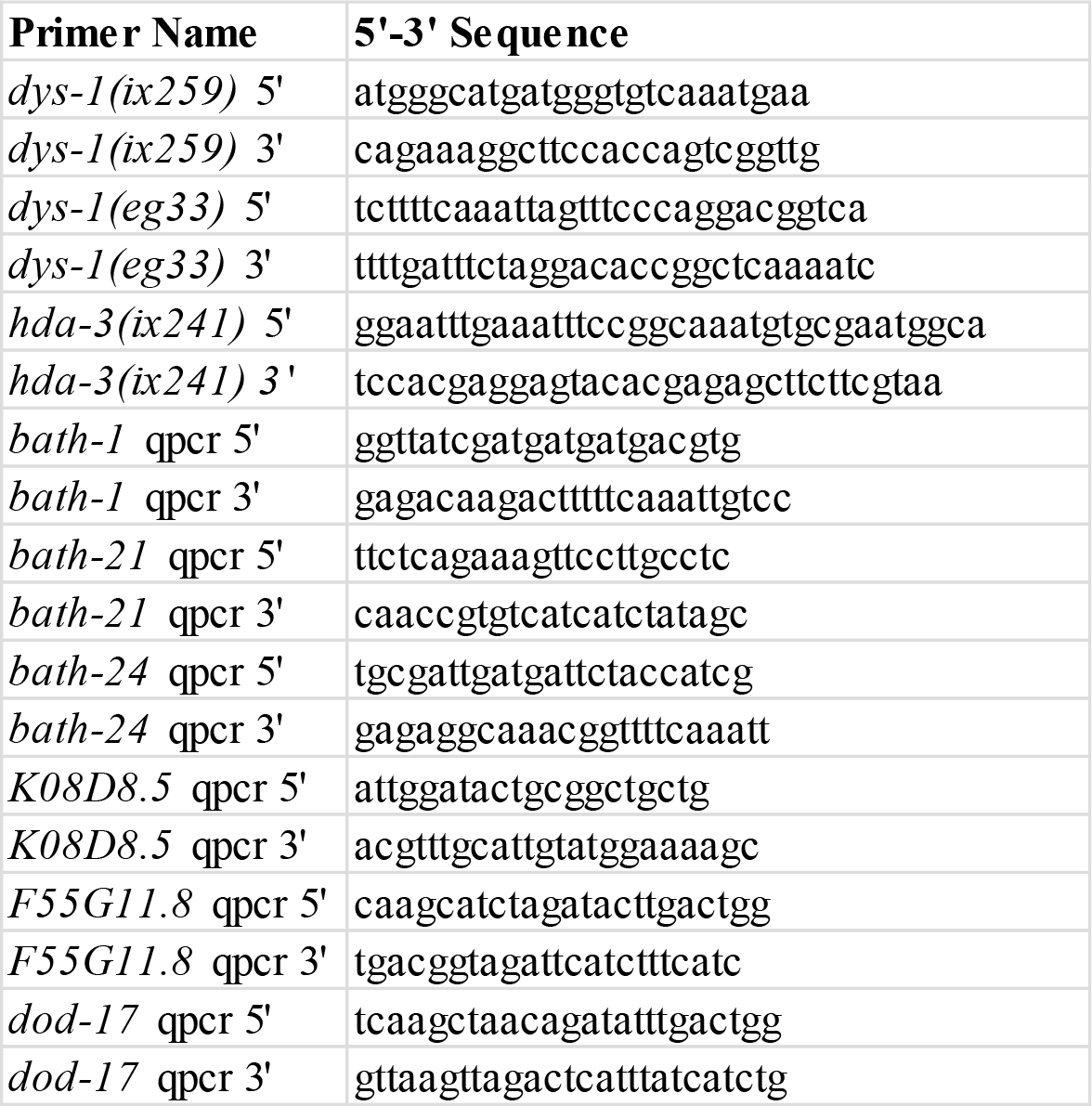
List of primers used in this study.

**Table S3.**
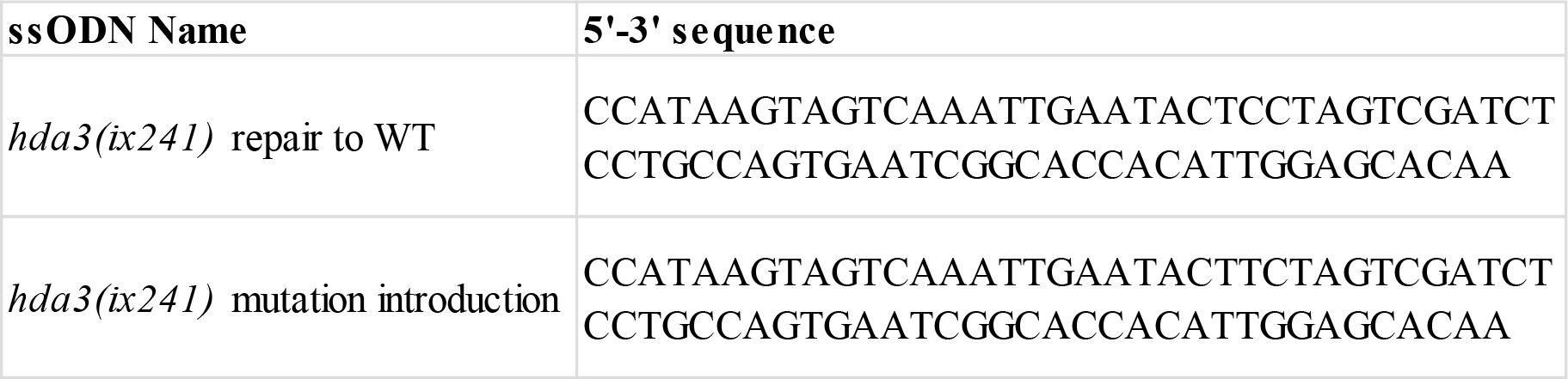
List of single-stranded oligodeoxynucleotide sequences used in this study.

**Table S4.**
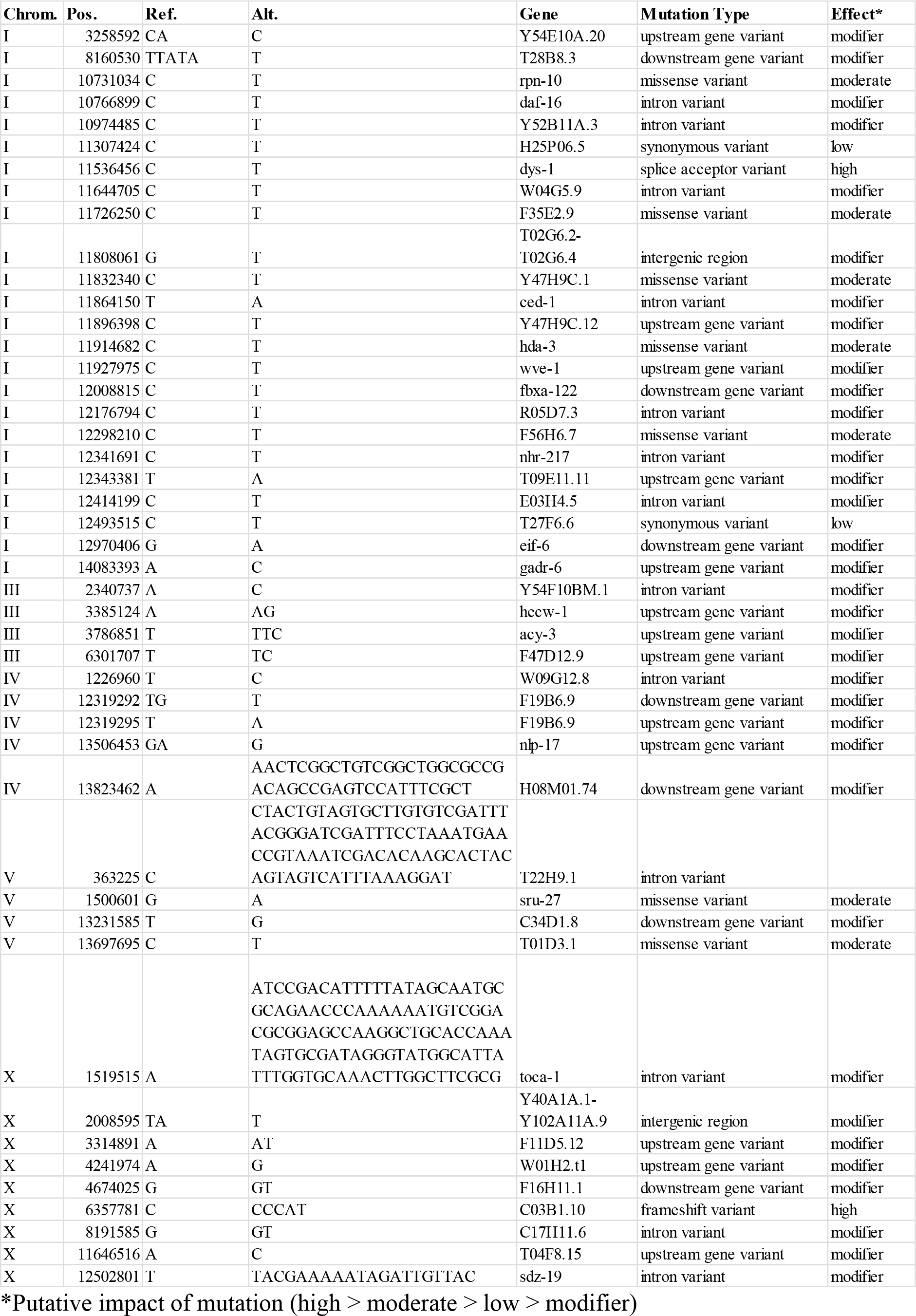
List of remaining mutations in backcrossed *ix241* strains.

**Table S5.** List of remaining mutations after subtraction of mutations in backcrossed *ix241* strains that do not show locomotor decline from *ix241* strains that show locomotor decline.

| Chrom. | Pos. | Ref. | Alt. | Gene | Mutation Type | Effect* |
| --- | --- | --- | --- | --- | --- | --- |
| I | 10731034 | C | T | rpn-10 | missense variant | moderate |
| I | 10766899 | C | T | daf-16 | intron variant | modifier |
| I | 10974485 | C | T | Y52B11A.3 | intron variant | modifier |
| I | 11307424 | C | T | H25P06.5 | synonymous variant | low |
| I | 11864150 | T | A | ced-1 | intron variant | modifier |
| I | 11896398 | C | T | Y47H9C.12 | upstream gene variant | modifier |
| I | 11914682 | C | T | hda-3 | missense variant | moderate |
| I | 11927975 | C | T | wve-1 | upstream gene variant | modifier |
| I | 12008815 | C | T | fbxa-122 | downstream gene variant | modifier |
| I | 12176794 | C | T | R05D7.3 | intron variant | modifier |
| I | 12298210 | C | T | F56H6.7 | missense variant | moderate |
| I | 12341691 | C | T | nhr-217 | intron variant | modifier |
| I | 12343381 | T | A | T09E11.11 | upstream gene variant | modifier |
| I | 12414199 | C | T | E03H4.5 | intron variant | modifier |
| I | 12493515 | C | T | T27F6.6 | synonymous variant | low |
| I | 12970406 | G | A | eif-6 | downstream gene variant | modifier |

\*Putative impact of mutation (high > moderate > low > modifier)

## Literature Cited

Bento-Abreu A, Jager G, Swinnen B, Rué L, Hendrickx S, Jones A, Staats KA, Taes I, Eykens C, Nonneman A, et al. 2018. Elongator subunit 3 (ELP3) modifies ALS through tRNA modification. Hum Mol Genet 27: 1276–1289. doi:10.1093/hmg/ddy043.

Bessou C, Giugia JB, Franks CJ, Holden-Dye L, Ségalat L. 1998. Mutations in the Caenorhabditis elegans dystrophin-like gene dys-1 lead to hyperactivity and suggest a link with cholinergic transmission. Neurogenetics 2: 61–72. doi:10.1111/imr.12206.

Blankenberg D, Kuster G Von, Coraor N, Ananda G, Lazarus R, Mangan M, Nekrutenko A, Taylor J. 2010. Galaxy: A web-based genome analysis tool for experimentalists. Curr Protoc Mol Biol 1–21. doi:10.1002/0471142727.mb1910s89.

Bork P, Beckmann G. 1993. The CUB Domain. J Mol Biol 231: 539–545. doi:10.1006/jmbi.1993.1305.

Brenner S. 1974. The genetics of Caenorhabditis elegans. Genetics 77: 71–94.

Cingolani P, Platts A, Wang LL, Coon M, Nguyen T, Wang L, Land SJ, Lu X, Ruden DM. 2012. A program for annotating and predicting the effects of single nucleotide polymorphisms, SnpEff: SNPs in the genome of Drosophila melanogaster strain w 1118; iso-2; iso-3. Fly (Austin) 6: 80–92. doi:10.4161/fly.19695.

Dobin A, Davis CA, Schlesinger F, Drenkow J, Zaleski C, Jha S, Batut P, Chaisson M, Gingeras TR. 2013. STAR: Ultrafast universal RNA-seq aligner. Bioinformatics 29: 15–21. doi:10.1093/bioinformatics/bts635.

Dokshin GA, Ghanta KS, Piscopo KM, Mello CC. 2018. Robust genome editing with short single-stranded and long, partially single-stranded DNA donors in caenorhabditis elegans. Genetics 210: 781–787. doi:10.1534/genetics.118.301532.

Eberharter A, Becker PB. 2002. Histone acetylation: A switch between repressive and permissive chromatin. Second in review on chromatin dynamics. EMBO Rep 3: 224–229. doi:10.1093/embo-reports/kvf053.

Giardine B, Riemer C, Hardison RC, Burhans R, Elnitski L, Shah P, Zhang Y, Blankenberg D, Albert I, Taylor J, et al. 2005. Galaxy: A platform for interactive large-scale genome analysis. Genome Res 15: 1451–1455. doi:10.1101/gr.4086505.

Goecks J, Nekrutenko A, Taylor J. 2010. Galaxy: a comprehensive approach for supporting accessible, reproducible, and transparent computational research in the life sciences. Genome Biol 11: R86. doi:10.1186/gb-2010-11-8-r86.

Grisoni K, Gieseler K, Mariol MC, Martin E, Carre-Pierrat M, Moulder G, Barstead R, Ségalat L. 2003. The stn-1 syntrophin gene of C. elegans is functionally related to dystrophin and dystrobrevin. J Mol Biol 332: 1037–1046. doi:10.1016/j.jmb.2003.08.021.

Groessl EJ, Kaplan RM, Rejeski WJ, Katula JA, King AC, Frierson G, Glynn NW, Hsu FC, Walkup M, Pahor M. 2007. Health-Related Quality of Life in Older Adults at Risk for Disability. Am J Prev Med 33: 214–218. doi:10.1016/j.amepre.2007.04.031.

Haberland M, Montgomery RL, Olson EN. 2009. The many roles of histone deacetylases in development and physiology: implications for disease and therapy. Nat Rev Genet 10: 32–42. doi:10.1038/nrg2485.

Hewitt JE, Pollard AK, Lesanpezeshki L, Deane CS, Gaffney CJ, Etheridge T, Szewczyk NJ, Vanapalli SA. 2018. Muscle strength deficiency and mitochondrial dysfunction in a muscular dystrophy model of Caenorhabditis elegans and its functional response to drugs. Dis Model Mech 11: dmm036137. doi:10.1242/dmm.036137.

Hoffman EP, Brown RH, Kunkel LM. 1987. Dystrophin: The protein product of the duchenne muscular dystrophy locus. Cell 51: 919–928. doi:10.1016/0092-8674(87)90579-4.

Kawamura K, Maruyama IN. 2019. Forward Genetic Screen for Caenorhabditis elegans Mutants with a Shortened Locomotor Healthspan. G3 (Bethesda) g3.400241.2019. doi:10.1534/g3.119.400241.

Kim H, Pierce-Shimomura JT, Oh HJ, Johnson BE, Goodman MB, McIntire SL. 2009. The Dystrophin Complex Controls BK Channel Localization and Muscle Activity in Caenorhabditis elegans. PLoS Genet 5: e1000780. doi:10.1371/journal.pgen.1000780.

Kim H, Rogers MJ, Richmond JE, McIntire SL. 2004. SNF-6 is an acetylcholine transporter interacting with the dystrophin complex in Caenorhabditis elegans. Nature 430: 891–896. doi:10.1038/nature02798.

Koboldt DC, Chen K, Wylie T, Larson DE, McLellan MD, Mardis ER, Weinstock GM, Wilson RK, Ding L. 2009. VarScan: Variant detection in massively parallel sequencing of individual and pooled samples. Bioinformatics 25: 2283–2285. doi:10.1093/bioinformatics/btp373.

Li H, Durbin R. 2009. Fast and accurate short read alignment with Burrows-Wheeler transform. Bioinformatics 25: 1754–1760. doi:10.1093/bioinformatics/btp324.

Li H, Handsaker B, Wysoker A, Fennell T, Ruan J, Homer N, Marth G, Abecasis G, Durbin R. 2009. The Sequence Alignment/Map format and SAMtools. Bioinformatics 25: 2078–2079. doi:10.1093/bioinformatics/btp352.

Liao Y, Smyth GK, Shi W. 2014. FeatureCounts: An efficient general purpose program for assigning sequence reads to genomic features. Bioinformatics 30: 923–930. doi:10.1093/bioinformatics/btt656.

Millard CJ, Watson PJ, Celardo I, Gordiyenko Y, Cowley SM, Robinson C V., Fairall L, Schwabe JWR. 2013. Class I HDACs share a common mechanism of regulation by inositol phosphates. Mol Cell 51: 57–67. doi:10.1016/j.molcel.2013.05.020.

Minevich G, Park DS, Blankenberg D, Poole RJ, Hobert O. 2012. CloudMap: a cloud-based pipeline for analysis of mutant genome sequences. Genetics 192: 1249–69. doi:10.1534/genetics.112.144204.

Montgomery RL, Potthoff MJ, Haberland M, Qi X, Matsuzaki S, Humphries KM, Richardson JA, Bassel-Duby R, Olson EN. 2008. Maintenance of cardiac energy metabolism by histone deacetylase 3 in mice. J Clin Invest 118: 3588–3597. doi:10.1172/JCI35847.

Nusinzon I, Horvath CM. 2005. Histone deacetylases as transcriptional activators? Role reversal in inducible gene regulation. Sci STKE 2005: re11. doi:10.1126/stke.2962005re11.

Oh HJ, Abraham LS, Van Hengel J, Stove C, Proszynski TJ, Gevaert K, DiMario JX, Sanes JR, Van Roy F, Kim H. 2012. Interaction of α-catulin with dystrobrevin contributes to integrity of dystrophin complex in muscle. J Biol Chem 287: 21717–21728. doi:10.1074/jbc.M112.369496.

Robinson MD, McCarthy DJ, Smyth GK. 2009. edgeR: A Bioconductor package for differential expression analysis of digital gene expression data. Bioinformatics 26: 139–140. doi:10.1093/bioinformatics/btp616.

Schapira M. 2011. Structural Biology of Human Metal-Dependent Histone Deacetylases. In Histone Deacetylases: the Biology and Clinical Implication (eds. T.-P. Yao and E. Seto), pp. 225–240, Springer Berlin Heidelberg, Berlin, Heidelberg doi:10.1007/978-3-642-21631-2_10.

Schuetz A, Min J, Allali-Hassani A, Schapira M, Shuen M, Loppnau P, Mazitschek R, Kwiatkowski NP, Lewis TA, Maglathin RL, et al. 2008. Human HDAC7 Harbors a Class IIa Histone Deacetylase-specific Zinc Binding Motif and Cryptic Deacetylase Activity. J Biol Chem 283: 11355–11363. doi:10.1074/jbc.M707362200.

Shi Y, Mello C. 1998. A CBP/p300 homolog specifies multiple differentiation pathways in Caenorhabditis elegans. Genes Dev 12: 943–55. doi:10.1101/gad.12.7.943.

Simpson CL, Lemmens R, Miskiewicz K, Broom WJ, Hansen VK, van Vught PWJ, Landers JE, Sapp P, Van Den Bosch L, Knight J, et al. 2009. Variants of the elongator protein 3 (ELP3) gene are associated with motor neuron degeneration. Hum Mol Genet 18: 472–81. doi:10.1093/hmg/ddn375.

Struhl K. 1998. Histone acetylation and transcriptional regulatory mechanisms. Genes Dev 12: 599–606. doi:10.1101/gad.12.5.599.

Team RC. 2015. R: A language and environment for statistical computing [Internet]. Vienna, Austria: R Foundation for Statistical Computing; 2015.

Thomas JH. 2006. Adaptive evolution in two large families of ubiquitin-ligase adapters in nematodes and plants. Genome Res 16: 1017–1030. doi:10.1101/gr.5089806.

Trombetti A, Reid KF, Hars M, Herrmann FR, Pasha E, Phillips EM, Fielding RA. 2016. Age-associated declines in muscle mass, strength, power, and physical performance: impact on fear of falling and quality of life. Osteoporos Int 27: 463–471. doi:10.1007/s00198-015-3236-5.

Wang A, Kurdistani SK, Grunstein M. 2002. Requirement of Hos2 histone deacetylase for gene activity in yeast. Science 298: 1412–4. doi:10.1126/science.1077790.

Watson PJ, Fairall L, Santos GM, Schwabe JWR. 2012. Structure of HDAC3 bound to co-repressor and inositol tetraphosphate. Nature 481: 335–340. doi:10.1038/nature10728.

Zhou S, Chen L. 2011. Neural integrity is maintained by dystrophin in C. elegans. J Cell Biol 192: 349–363. doi:10.1083/jcb.201006109.

